# Transcript-wide m^6^A methylation defines the efficiency of the cap-independent translation initiation

**DOI:** 10.1101/2025.11.25.690160

**Authors:** Egor A. Smolin, Andrey I. Buyan, Anton A. Buzdin, Dmitry N. Lyabin, Ivan V. Kulakovskiy, Irina A. Eliseeva

## Abstract

N^6^-methyladenosine (m^6^A) role in translation control and, particularly, in cap-independent initiation has attracted major attention. A common approach is to study the impact of m^6^A depending on its particular location in the transcripts, but the global impact of m^6^A along mRNAs remains unclear.

Here, we combined ribosome profiling under conditions of mTOR inhibition and m^6^A mapping to identify distinct subsets of mRNAs that differ in their sensitivity to the suppression of cap-dependent initiation. The sensitivity was strongly correlated with the total m^6^A methylation of the transcripts. Further, we demonstrated that upon decreased mTOR activity, m^6^A methylation facilitated the enhanced association of mRNAs with components of the eIF4F complex.

Thus, the efficiency of cap-independent translation initiation is primarily defined not by precise localization but by the total level of m^6^A methylation, and m^6^A has a compensatory role in maintaining translation when the canonical cap-dependent pathway is impaired. All in all, our findings underscore the significance of contemplating global m^6^A methylation status as a pivotal element in translational control, particularly under stress or signaling perturbations.

**GRAPHICAL ABSTRACT:** 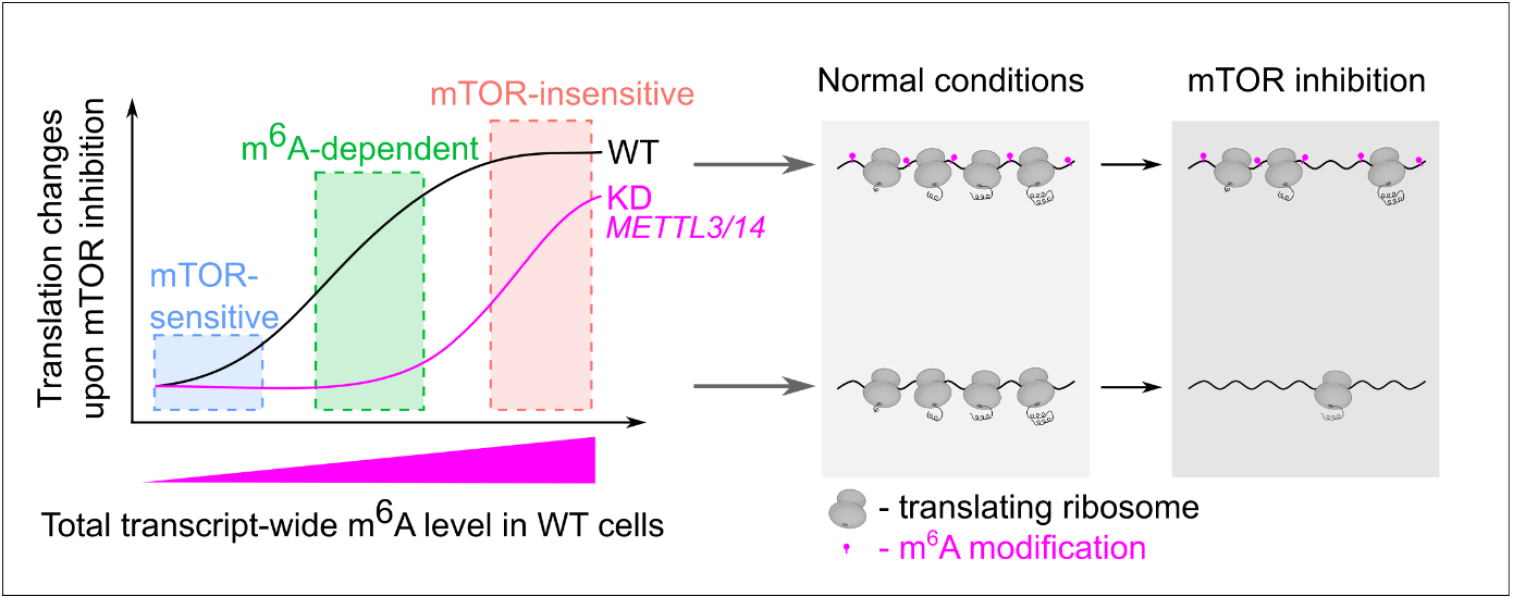

## INTRODUCTION

N^6^-methyladenosine (m^6^A) is the most abundant epitranscriptomic modification of eukaryotic mRNAs. It has been implicated in diverse cellular processes, including mRNA splicing, nuclear export, decay, and translation^1^. m^6^A influence on translation control has been actively studied in the last decade^2^, but its role in cap-independent translation has not been explored in full.

Under normal conditions, translation initiation is typically mediated by the recruitment of the 43S translation pre-initiation complex to mRNA via the interaction of the eIF4F complex (composed of eIF4E, eIF4G, and eIF4A) with the 5′ cap structure and adjacent sequences. Under stress conditions, translation can initiate independently of the cap structure and the cap-binding initiation factor eIF4E^3–5^. The central pathway regulating eIF4E activity is the mTOR signaling pathway, which responds to cellular stress, nutrient availability, and energy^6,7^. One of its key mTOR targets is the eIF4E-binding protein 1 (4E-BP1), which, in its hypophosphorylated state, binds eIF4E and inhibits the assembly of the eIF4F complex, thereby suppressing cap-dependent translation. Activation of mTOR leads to the hyperphosphorylation of 4E-BP1, which releases eIF4E and restores cap-dependent translation initiation^8–10^.

Notably, the canonical mTOR-sensitive mRNAs are largely devoid of m^6^A modifications^11^. Furthermore, knockdown of *METTL3*, the catalytic subunit of the m^6^A methyltransferase complex, increases the sensitivity of global protein synthesis to mTOR inhibition^12^. These findings suggest that m^6^A may contribute to cap-independent translational control, probably supporting a compensatory mechanism when cap-dependent translation is compromised.

High-throughput mapping revealed that m^6^A is predominantly enriched within the 3′ untranslated regions (3′ UTRs) and, specifically, in the vicinity of the stop codons, with lower levels in the 5′ UTRs and coding segments^13,14^. When m^6^A is located in 5′ UTRs, it can promote cap-independent translation initiation by facilitating eIF3 or ABCF1 binding to the mRNA. In turn, these proteins recruit the 43S pre-initiation complex to the mRNA^11,12^. This phenomenon is of particular interest within the context of mTOR-dependent translational regulation as canonical mTOR target mRNAs lack m^6^A, particularly within 5’ UTRs^12^. This may explain why these mRNAs are more sensitive to eIF4E inactivation.

Yet, contrary to previous assertions, recently it has been demonstrated that the presence of m^6^A exclusively in 5’ UTRs does not significantly enhance translation efficiency or promote the assembly of the pre-initiation complex^15^. Furthermore, 5′ UTRs in general carry fewer m^6^As compared to the other mRNA regions. In 3’ UTRs, the interactions between m^6^A and the “readers” such as YTHDF1 or METTL3 can facilitate the recruitment of eIF3, thereby promoting translation^16,17^. Furthermore, m^6^A in 3′ UTR contributes to mRNA circularization through interactions with YTHDF3, enhancing the efficiency of translation re-initiation^18^.

When m^6^A is localized in coding regions, it can modulate the translation elongation rate. Depending on the context, m^6^A can either accelerate^19^ or slow down elongation^20^, probably by affecting the accuracy of codon-anticodon recognition^21^. Particularly, m^6^A hinders the tRNA interaction with the ribosome, provoking premature detachment and enhanced proofreading, which reduces translation efficiency^22^. Further, m^6^A in the coding regions can trigger translation-dependent mRNA decay, which is mainly active at low ribosome load or under cellular stress^23^.

All in all, the generally accepted concept is that the effect of m^6^A on translation critically depends on its position within the transcript. We challenge this concept by showing that mRNA sensitivity to mTOR inhibition is associated with the overall level of m^6^A methylation along the transcript. These findings imply that the presence of m^6^A, rather than its precise position within the transcript, is critical for sustaining translation under mTOR inhibition. Further, we show that in these conditions, mRNAs with the elevated m^6^A levels exhibit augmented association with the eIF4F complex.

## MATERIALS AND METHODS

### Plasmids

The 3′ untranslated regions (UTRs) of *SLU7* and *RPL32* mRNAs, as well as the promoter region, 5′ UTR, and 3′ UTR of the *HSPA1A* mRNA (*HSP70*), were amplified from HEK293T genomic DNA using the Phusion High-Fidelity PCR Kit (Thermo Fisher Scientific, Waltham, MA, USA) in a GC-rich buffer. The primers included 20-nucleotide overhangs for ligation-independent cloning (SLIC) (see **Table S1**).

The purified PCR fragments containing the 3′ UTRs of *SLU7* mRNA and *RPL32* mRNA were cloned into plasmids pNL2.2 *SLU7-NlucP* and pNL2.2 *RPL32-NlucP*^24^, respectively, using SLIC (primers used for vector preparation are listed in **Table S1**). The promoter region, 5′ UTR, and 3′ UTR of the *HSPA1A* mRNA were cloned into a promoterless pNL2.2 vector (Promega, Madison, WI, USA) using the same method.

The verified plasmids were named as follows: pNL2.2 *SLU7-NlucP-SLU7*, pNL2.2 *RPL32-NlucP-RPL32*, and pNL2.2 *HSPA1A-NlucP-HSPA1A*.

The pcDNA3.1-HA was constructed previously^25^.

### Cell cultures

HEK293T cells (originally obtained from ATCC) were provided by Dr. Elena Nadezhdina (Institute of Protein Research, RAS). The cells were cultured in Dulbecco’s Modified Eagle Medium (DMEM, Capricorn Scientific, Germany) supplemented with 10% fetal bovine serum, 2 mM glutamine, 100 U/mL penicillin, and 100 μg/mL streptomycin. Cultures were maintained at 37 °C in a humidified atmosphere containing 5% CO_2_ and passaged using standard procedures. To inhibit mTOR kinase, cells were treated with Torin1 (250 nM; Tocris Bioscience, Bristol, UK), dissolved in DMSO, or with DMSO alone as a vehicle control. Treatment times are indicated in the figure legends.

### *METTL3/14* knockdown with siRNA

To investigate the role of endogenous m^6^A methylation, we used wild-type HEK293T cells transfected with either a non-targeting control siRNA or a mixture of siRNAs targeting *METTL3* and *METTL14* (see **Table S1**). To anneal siRNA duplexes, an equal molar mixture of individual RNA strands in an annealing buffer (40 mM Tris-HCl, 10 mM MgCl_2_, 10 mM DTT, pH 7.8) was heated at 95°C for 5 minutes, then slowly cooled to 37°C over 30-60 minutes.

siRNA transfection was performed using HiPerFect reagent (Qiagen, Germany) following the manufacturer’s reverse transfection protocol. Briefly, cells were seeded at approximately 50% confluency just before adding the siRNA-HiPerFect complexes. For each 100 mm dish, 10 nM siRNA with 30 μL of HiPerFect in 200 μL of Opti-MEM was used.

To enhance knockdown efficiency, a second transfection was conducted 24 hours later. Cells were collected for downstream experiments 48 hours after the initial transfection. The knockdown efficiency was validated by Western blot analysis.

### DNA and mRNA transfection

*For DNA transfections*, HEK293T cells were transfected using Lipofectamine 3000 (Thermo Fisher Scientific, Waltham, MA, USA) according to the manufacturer’s protocol. For each 6-well, 400 ng of pNL2.2 plasmid and 100 ng of pcDNA3.1-HA were combined with 1 μL of P3000 reagent and 2 μL of Lipofectamine 3000 in 100 μL of Opti-MEM (Gibco). After a 15-minute incubation at room temperature, the mixture was added to the cells. Following a 4–6 hour incubation, the cells were distributed into 48-well plates and further cultured under standard conditions for an additional 16 hours before treatment with Torin1.

*For mRNA transfection, in vitro* transcribed K^+^A^+^ reporter mRNAs were delivered using Lipofectamine 3000 in HEK293T that had been pretreated with Torin 1 for 2 hours. For each 48-well (**“m**^**6**^**A methylation of mRNAs affects their cap-independent translation initiation in HEK293” in results**), 300 ng of mRNA was diluted in 50 μL of Opti-MEM and mixed with 0.6 μL of Lipofectamine 3000. For each 96-well (**“The sensitivity to mTOR inhibition saturates at a certain m**^**6**^**A methylation level” in results**), 100 ng of mRNA was diluted in 10 μL of Opti-MEM, combined with 0.3 μL of Lipofectamine 3000, and supplemented with a 1 μL concentrated furimazine solution (Nano-Glo, Promega). In both cases, the mixture was incubated at room temperature for 15 minutes before being added to the cells.

*NanoLuc luciferase activity (NlucP)* was measured using the Nano-Glo Luciferase Assay System (Promega). Cells were lysed in Passive Lysis Buffer (PLB) (Promega) for 15 minutes at 37°C, and luminescence was measured using a GloMax 20/20 Luminometer (Promega). Real-time luminescence (**“The sensitivity to mTOR inhibition saturates at a certain m**^**6**^**A methylation level” in results**) was monitored immediately after transfection using CLARIOstar Plus (BMG LABTECH, Germany).

### Western blotting

For Western blot analysis, the cells were rinsed two times with phosphate-buffered saline (PBS) and lysed in SDS electrophoresis sample buffer. Proteins were separated by SDS-PAGE and transferred onto a nitrocellulose membrane (Cytiva, USA). The membrane was blocked for 1 h at room temperature with 5% BSA (Dia-M, Moscow, Russia) in TBS (10 mM Tris-HCl, pH 7.6, 150 mM NaCl) and then incubated overnight at 4°C in TBS-T (TBS with 0.05% Tween-20) supplemented with 5% BSA and the indicated primary antibodies (all from Cell Signaling Technology): phospho-p70 S6 kinase T389 (108D2), phospho-S6 ribosomal protein S235/236 (D57.2.2E), S6 ribosomal protein (5G10), phospho-4EBP1 T37/46 (236B4), 4EBP1 (53H11), METTL3 (D2I6O), METTL14 (D8K8W), mTOR (7C10), phospho-mTOR S2448 (49F9), AKT (C67E7), phospho-AKT S473 (D9E), eIF4E (C46H6), phospho-eIF4E S209 (F4E5N), eIF4A (F52), eIF4G (C45A4), β-Actin (13E5) and Fibrillarin (C13C3). Fibrilarin antibodies were used at 1:10000 dilution, and other primary antibodies were used at 1:2000 dilution.

After washing three times with TBS-T, the membrane was incubated for 1 h at room temperature with horseradish peroxidase (HRP)-conjugated goat anti-rabbit IgG secondary antibody (1:4,000, #7074, Cell Signaling Technology) diluted in 5% BSA in TBS-T. Following three additional washes with TBS-T, the immunocomplexes were detected using the ECL Prime kit (Cytiva, USA) according to the manufacturer’s instructions.

### RNA isolation

Cells were collected in ice-cold PBS, and total RNA was extracted using QIAzol Lysis Reagent (Qiagen, Germany) and Direct-Zol RNA Microprep kit (Zymo Research, USA) according to the manufacturer’s recommendations.

### *In vitro* transcription and RNA modification

For *in vitro* transcription, the PCR product generated with gene-specific primers that contain an encoded T7 promoter (**Table S1**) was used as a template. *In vitro* transcription was performed using the T7-Scribe Standard RNA IVT Kit (CellScript, USA) according to the manufacturer’s instructions. m^6^A-Methylated mRNAs were generated by incorporating m^6^ATP (Jena Bioscience, Germany, NU-1101L) along with ATP at defined ratios.

Capping and polyadenylation were conducted using the ScriptCap™ m7G Capping System and the A-Plus™ Poly(A) Polymerase Tailing Kit (CellScript, USA) following the manufacturer’s protocols.

For RNA pull-down experiments, after transcription, mRNAs were initially polyadenylated, then 3’-biotinylated as previously described^26^, and subsequently enzymatically capped using the same kits mentioned above.

RNA quality was evaluated by electrophoresis in a 1% agarose gel containing 2.2 M formaldehyde.

### m^6^A detection

m^6^A levels were evaluated by dot blot analysis. RNA samples were denatured at 70 °C for 5 min, cooled on ice, and mixed with an equal volume of cold 10× SSC buffer (1.5 M NaCl and 150 mM sodium citrate, pH 7.0)^27^. Samples were applied to Amersham Hybond-N^+^ membranes (Cytiva, USA) using a Bio-Dot apparatus (Bio-Rad). Membranes were UV-crosslinked (1200 μJ/cm^2^; Vilber Lourmat FLX-20M), rinsed briefly with water, and stained with 0.02% methylene blue in 0.3 M sodium acetate (pH 5.2) for 5–10 min to verify equal RNA loading. After destaining with water, membranes were air-dried and then blocked in 7.5% BSA in PBST (PBS with 0.05% Tween-20) for 60 min at room temperature. Membranes were incubated overnight at 4 °C with rabbit anti-m^6^A antibody (1:3000; Abcam, #151230) in blocking buffer, washed three times with PBST, and incubated for 1 h with HRP-conjugated goat anti-rabbit IgG (1:5000; Cell Signaling Technology, #7074) in 7.5% BSA/PBST. After three additional washes with PBST, immunocomplexes were detected using the ECL Prime detection system (Cytiva, USA) according to the manufacturer’s instructions.

### Cell-free translational system (CFTS)

The cell extract-based cell-free translation system was obtained, as previously described^28^, from HEK293T cells treated with 250nM Torin1 or DMSO for 2 h.

The translation reaction (10 μL) contained 5 μL HEK293T extract, 1 μL 10× Translation buffer (200 mM HEPES-KOH, pH 7.6, 10 mM DTT, 5 mM spermidine-HCl, 80 mM creatine phosphate, 10 mM ATP, 2 mM GTP, and 250 μM of each amino acid), 100 mM KOAc, 1 mM Mg(OAc)_2_, 2 U of Human Placental RNase Inhibitor (Thermo Fisher Scientific, USA), and 0.15 pmol of reporter *NlucP* mRNA. Reactions were incubated at 30 °C for 40 min, and luciferase activity was measured using the Nano-Glo Luciferase Assay System (Promega, USA) according to the manufacturer’s instructions.

Where indicated, m^7^GpppG or ApppG cap analogs were added at 0.1 mM concentrations. In such cases, translation mixtures were preincubated for 5 min at 30 °C prior to the addition of mRNA.

For real-time luminescence (**“The sensitivity to mTOR inhibition saturates at a certain m**^**6**^**A methylation level” in results**), the CFTS was supplemented with 1 µL of concentrated furimazine solution (Nano-Glo, Promega) to enable continuous monitoring of luciferase activity in real time. Real-time luminescence was recorded immediately after transfection using a CLARIOstar Plus plate reader (BMG LABTECH, Germany).

### RNA pull-down assay

Streptavidin-Sepharose (Cytiva, USA) was washed five times with 30 volumes of PBS supplemented with 0.2 mM VRC (Fluka, Switzerland). Then the resin was incubated overnight with a blocking solution (PBS supplemented with 1 mg/ml BSA, 200 μg/ml glycogen, and 200 μg/ml total RNA from *Escherichia coli*) at 4 °C with constant stirring. After incubation, the resin was washed five times with 10 volumes of PBS and 2 times with 10 volumes of for Translation buffer (20 mM Hepes–KOH, pH 7.6, 1 mM DTT, 0.5 mM spermidine–HCl, 1 mM Mg(OAc)_2_, 8 mM creatine phosphate, 1 mM ATP, 0.2 mM GTP, 100 mM KOAc and 25 μM of each amino acid). The 2 pmol capped biotinylated RNA was incubated with 50 μl HEK293T cell extract (for CFTS) in Translation buffer supplemented with 20 μg of total *E*.*coli* RNA for 15 min at 30 °C. This mixture was then supplemented with 35 μl of 50% pre-blocked Streptavidin-Sepharose slurry and incubated for 1 h at room temperature. The resin with RNA–protein complexes was washed 8 times with 30 volumes of PBS (with 0.2 mM VRC). Bound proteins were eluted with the SDS-PAGE sample buffer (80 mM Tris-HCl, pH 6.8, 2% SDS, 200 mM 2-mercaptoethanol, 10% glycerol, and 0.0012% bromophenol blue), resolved by 10% SDS-PAGE, and analyzed by Western blotting.

### Ribo-Seq and RNA-Seq experiments

HEK293T (WT) and HEK293T *METTL3/METTL14* knockdown (KD) cells were cultured in the absence or presence of Torin1 (250 nM) until they reached 70–80% confluency. The cells were immediately chilled on ice and washed with PBS supplemented with cycloheximide (100 µg/mL). The cells were lysed directly on the dish using a buffer containing 20 mM Tris-HCl (pH 7.4), 150 mM NaCl, 5 mM MgCl_2_, 1 mM DTT, 1% Triton X-100, 100 µg/mL cycloheximide (Sigma-Aldrich, USA), and 25 U/mL TURBO DNase (Ambion, Thermo Fisher Scientific, USA). Cell lysates were incubated on ice for 10 min, triturated five times through a 26-G needle, and centrifuged at 20,000×g at 4 °C for 10 min. The supernatant was divided into two parts for Ribo-seq and RNA-seq library preparation. Nuclease footprinting and ribosome recovery for Ribo-seq library preparation were performed according to the protocol described in ^29^. The adapter used for the Ribo-seq library (**Table S1**) contained a UMI sequence (five random nucleotides) to allow for the deduplication of the resulting reads (see below). Subsequent procedures for Ribo-Seq were performed as described in^29^.

Total RNA for RNA-Seq was isolated using TRIzol LS Reagent (Thermo Fisher Scientific, USA) and Direct-Zol RNA Miniprep kit (Zymo Research, USA). rRNA depletion was performed using a RiboMinus™ Eukaryote Kit v2 (Thermo Fisher Scientific, USA) according to the manufacturer’s instructions. RNA-Seq libraries were generated using a NEBNext® Ultra™ II Directional RNA Library Prep Kit for Illumina (New England Biolabs, USA) following the manufacturer’s protocol. For the read counts normalization in the consequence data analysis, spike-in RNAs of known concentration and unrelated to the human genome (*PSP73, GFP*, and Ribo spike-in), generated by *in vitro* transcription, were added to each sample during library preparation (**Table S2**).

### High-Throughput Sequencing and Data Processing

#### Sequencing and basic processing

The NGS libraries were sequenced on the Illumina platforms (RNA-Seq: Illumina HiSeq 2500, 60bp single-end reads; Ribo-Seq: Illumina HiSeq 4000, 51bp single-end reads), yielding 17 mln. reads per sample on average. Read quality control was performed with FastQC v0.11.5. The reads were trimmed with cutadapt v2.10 ^30^: “*--minimum-length 20 -q 20” for RNA-Seq and “-a ATCGTAGATCGGAAGAGCACACGTCTGAA -j 24 --minimum-length 20 --nextseq-trim=20 --trimmed-only*” for Ribo-Seq. For the latter, the availability of UMIs allowed pre-alignment deduplication using seqkit v2.6.1 rmdup ^31^ followed by cutadapt “*-u -5*” to remove UMIs.

#### Read mapping and additional quality control

Read mapping was performed to the hg38 genome assembly using STAR v2.7.6a^32^ with default parameters and “*--quantMode GeneCounts*” to estimate the read counts per gene (**Table S2**). The genome annotation was based on GENCODE v38 and modified to include spike-in sequences as additional contigs. Specifically for Ribo-Seq, the quality check was performed using plastid (v.0.5.1) *generate*^33^, *psite*, and *phase_by_size* commands, following P-site offset identification and read phasing (**Figure S1A**) protocols as described in the plastid documentation. The agreement of replicates was qualitatively assessed by PCA (**Figure S1B**).

#### Batch correction and differential expression analysis

RNA-Seq and Ribo-Seq gene count matrices were filtered, leaving only spike-ins and protein-coding genes encoded in the nuclear genome with at least 5 cpm (for RNA-Seq, the minimal library size of 2217774) or 15 cpm (for Ribo-Seq, the minimal library size of 735116) in more than 2 samples. Batch correction was performed using *ComBat-seq* from sva (v.3.35.2) and WT^34^, WT + Torin, KD, and KD + Torin cells as biological groups. Differential expression and ribosome occupancy (RNA-Seq-normalized ribosome footprint density as a proxy for the translation efficiency) were calculated with DESeq2 (v1.38.1)^35^. Differential expression analysis was performed with DESeq2 *DESeqDataSetFromMatrix, estimateSizeFactors, estimateDispersions*, and *nbinomLRT* commands using *PSP73* and *GFP* spike-ins (for RNA-Seq) and Ribo spike-in (for Ribo-Seq) as control genes.

#### Comparison of differential ribosome occupancy for KD and WT cells

Differential ribosome occupancy analysis resulted in several gene sets: downregulated under stress conditions both in WT and KD cells (mTOR-sensitive, 3849 genes with *WT_logFC < -0*.*6 & WT_padj < 0*.*05 & DD_logFC < -0*.*6 & DD_padj < 0*.*05*), downregulated only in KD cells (m^6^A-dependent, 569 genes with, *(WT_logFC > -0*.*4 & WT_padj > 0*.*05* | *WT_logFC > 0 & WT_padj < 0*.*05) & DD_logFC < -0*.*6 & DD_padj < 0*.*05*) and unchanged under stress conditions in both cells (mTOR-insensitive, 704 genes with *abs(WT_logFC) < 0*.*3 & WT_padj > 0*.*05 & abs(DD_logFC) < 0*.*3 & DD_padj > 0*.*05*). To annotate these gene groups, we have performed hypergeometric tests for KEGG and GO annotations using *kegga* and *goana* functions from limma (v.3.54.0)^36^ with org.Hs.eg.db (v.3.16.0)^37^ and *mapIds* from AnnotationDbi (v.1.60.0)^38^ being used to translate ENSEMBL gene ids into Entrez ids.

To examine sequence properties of transcripts belonging to these groups, we assigned the most expressed protein-coding transcript to each gene. Isoforms abundances were estimated using Salmon (v.0.14.1)^39^ *index* and *quant* commands, and GENCODE comprehensive annotation (v.38)^40^, yielding TPM values that were used to prioritize highly expressed transcripts. Finally, 5’ UTR, CDS, and 3’ UTR lengths and their CG-content (**Figure S2A**,**B**) were estimated for each transcript using BSgenome.Hsapiens.UCSC.hg38 (v.1.4.4) and *getSeq* command from BSgenome (v.1.66.1).

#### RBP targets enrichment analysis

We used one-tailed Fisher’s exact test for the association between each gene group and TOP genes^41^, eCLIP K562 and HepG2 targets^42^, and LARP1, YTHDF1, YTHDF2, WTAP, METTL14, and METTL3 targets^43–46^ (**Table S3**).

#### Analysis of m^6^A modifications

The existing data on m^6^A modifications were mapped to the prevalent transcripts and specifically their UTR and CDS regions using the most deeply sequenced replicate for eTAM-Seq^47^ and m^6^A sites identified in HEK293T cells for GLORI^48^. Genomic positions were converted into BED format and intersected with GENCODE v38 annotation (UTR and CDS features) using bedtools (v2.31.0)^49^ to locate m^6^A sites within individual transcripts and transcript regions (5′ UTR, CDS, 3′ UTR). The normalized total transcript m^6^A methylation level was estimated as the sum of the per-site m^6^A percentages along a transcript relative to the number of adenosines in the transcript.

## RESULTS

### Translation of m^6^A-modified mRNAs is less sensitive to mTOR inhibition

To investigate how m^6^A affects cap-independent initiation, we used *in vitro* transcribed reporter mRNAs with 5’ and 3’ UTRs of *RPL32, SLU7*, and *HSP70* (*HSPA1A*). *RPL32* is a classic mTOR mRNA target^50^, and *SLU7* served as the negative control unaffected by mTOR suppression^24^. *HSP70* mRNA belongs to the group of mRNAs for which methylation and translation efficiency increase under heat shock conditions when cap-dependent initiation is suppressed^51^.

All reporter mRNAs were methylated during *in vitro* transcription by incorporating m^6^A-ATP in a 1:1 ratio with ATP, ensuring a high level of m^6^A inclusion in the resulting transcripts. To inhibit cap-dependent initiation, we used a cell-free translation system prepared from HEK293T cells and two strategies: (1) the addition of a cap analog to compete with the native cap for eIF4E binding, and (2) treatment with Torin1, the widely used mTOR inhibitor.

m^6^A-modified mRNAs demonstrated the reduced sensitivity to translation inhibition in both cases (**Figure 1, A-B**), irrespective of their 5′ and 3′ UTRs. One could expect the effect to be UTR-specific^8,10^, reflecting the cellular differences between mRNAs in their mTOR sensitivity. Yet, in this case, the strong presence of m^6^A along the transcript was sufficient to promote resistance to cap-dependent inhibition.

**Figure 1.**
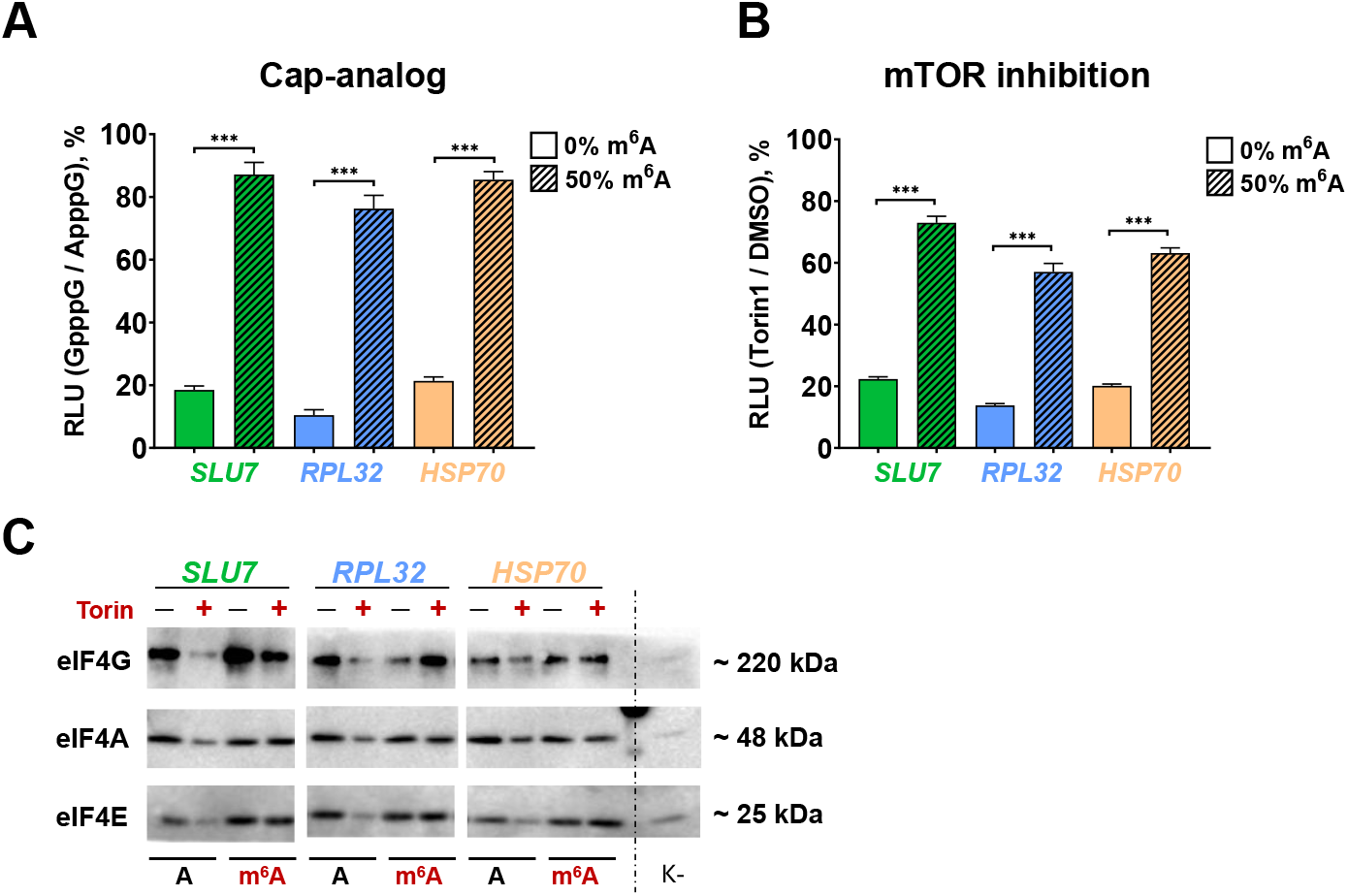
m^6^A enhances cap-independent translation *in vitro*. The *NlucP* reporter mRNAs with 5’ and 3’ UTRs of *SLU7, RPL32*, and *HSP70* were transcribed *in vitro* in the presence of 50% or 0% m^6^ATP. mRNAs were then enzymatically capped and polyadenylated **(A-B)**. For the pull-down assay **(C)**, after transcription, the mRNAs were first polyadenylated, then biotinylated at the 3’ end, and enzymatically capped. **(A)** mRNAs were translated in CFTS prepared from untreated HEK293T cells in the presence of mM cap-analogs m7GpppG or ApppG. NlucP activity in the presence of m7GpppG was normalized to that in the presence of ApppG. **(B)** mRNAs were translated in CFTS prepared from HEK293T cells treated with 250nM Torin1 or DMSO. NlucP activity in Torin1-treated CFTS was normalized to that in DMSO-treated CFTS. Values are the means of at least three independent replicates. Error bars show standard deviations. Two-tailed Student’s t-test was used to estimate the statistical significance between methylated and unmethylated mRNAs. ***p<0.001. **(C)** The biotinylated mRNAs were translated in CFTS prepared from HEK293T cells treated with 250nM Torin1 or DMSO. RNA-protein complexes were isolated using streptavidin-sepharose and analysed by Western blot using antibodies against eIF4A, eIF4E, and eIF4G. Uncapped mRNA with *RPL32* UTRs (K-) was used as a control. CFTS: cell-free translation system; RLU: relative luciferase units.

Interestingly, the impact of m^6^A was more exhibited in the case of the cap analog (4-7-fold, **Figure 1A**) compared to Torin-treated lysates (3-4-fold, **Figure 1B**). We interpret this difference as follows. The cap analog directly affects eIF4E availability and thus provides a targeted instrument to assess the contribution of m^6^A to the cap-dependent initiation efficiency. In turn, inhibition of mTOR signaling has many extra consequences, such as reduced eIF4A helicase activity through S6K-dependent mechanisms^52^, leading to reduced initiation efficiency of the tested m^6^A-modified mRNAs.

It is known that m^6^A-containing transcripts can interact with eIF3 or ABCF1, facilitating recruitment of the 43S pre-initiation complex^11,12^. Yet, it was unexplored whether m^6^A methylation affects the binding of the core cap-binding complex, eIF4F.

We performed pull-down assays to examine how the components of the eIF4F complex bind m^6^A-modified and unmethylated mRNAs under normal conditions and upon mTOR inhibition. Strikingly, eIF4A, eIF4G, and eIF4E have a higher affinity to m^6^A-modified mRNAs compared to their unmethylated counterparts (**Figure 1C**). These interactions remained stable even upon treatment with Torin1, suggesting that m^6^A-modified transcripts retain their ability to interact with eIF4F (**Figure 1C**).

Overall, m^6^A can enhance translation initiation by increasing the binding affinity for eIF4F when cap-dependent initiation is inhibited.

### m^6^A methylation of mRNAs affects their cap-independent translation initiation in HEK293

As shown above, in the cell-free translation system, m^6^A methylation reduced the sensitivity of mRNAs to the inhibition of cap-dependent initiation. However, in these assays, m^6^A was introduced in random locations along the entire length of the transcripts, which significantly differs from the endogenous distribution of m^6^A.

To evaluate the effect of m^6^A modifications natively distributed along transcripts on translation upon mTOR inhibition, we employed a reporter system, featuring 5’ UTRs and native promoters of *SLU7, RPL32*, and *HSP70* (**Figure 2B**). Here, were employed native promoters as the extent and localization of m^6^A in mRNAs depend on transcription initiation due to co-transcriptional recruitment of the methyltransferase complex^20^.

**Figure 2.**
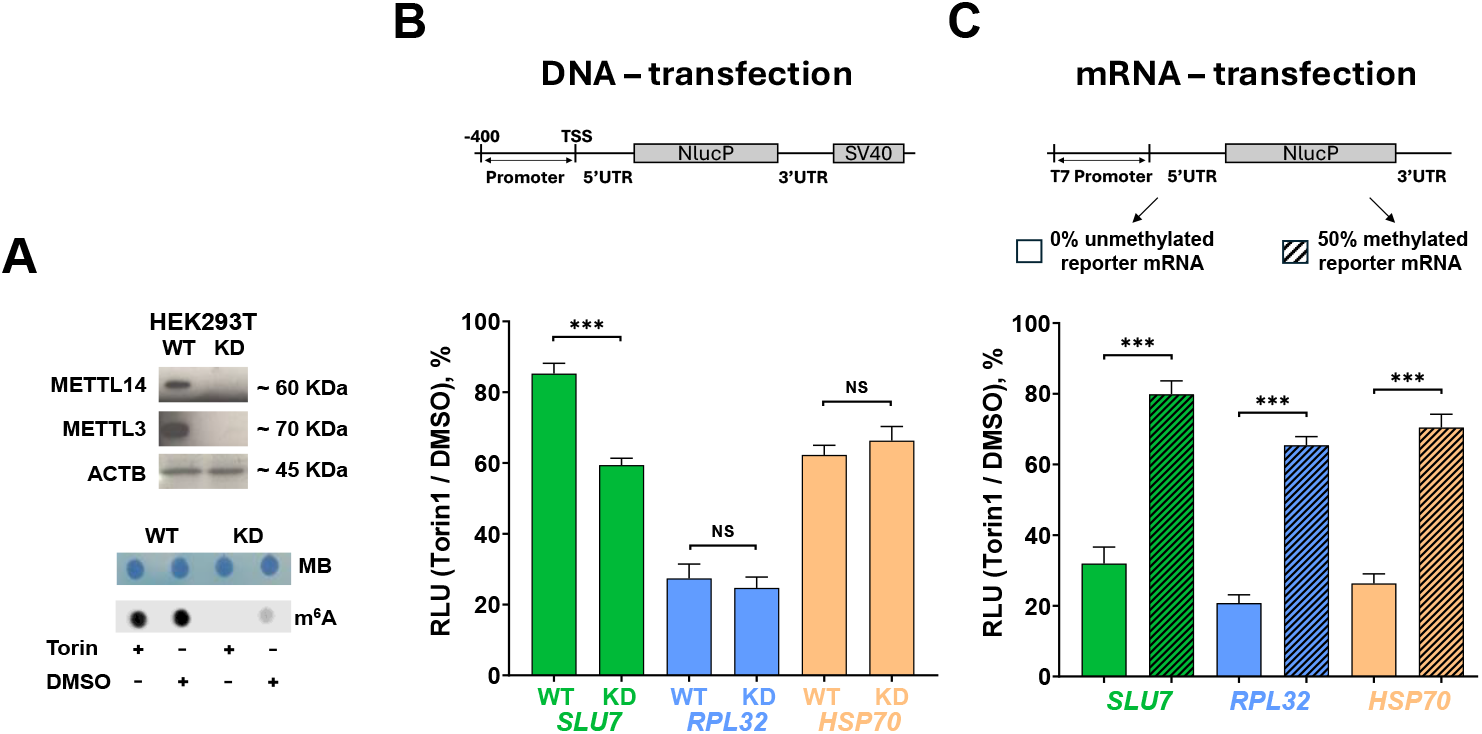
Lack of m^6^A impairs cap-dependent translation of mRNAs in response to mTOR inhibition in HEK293T cells. **(A)** *METTL3* and *METTL14* knockdown (KD) with siRNA transfection in HEK293T cells reduces m^6^A methylation level. **(top panel)** Western blot analysis of METTL3 and METTL14 abundance in WT and KD HEK293T cells. β-actin (ACTB) was used as a loading control. **(bottom panel)** Dot blot analysis of m^6^A methylation in total RNA from WT and KD HEK293T cells treated with 250 nM Torin1 or DMSO. Total RNA was transferred onto a nylon membrane and stained with antibodies against m^6^A or with methylene blue (MB). **(B)** Reporter DNA vectors were transfected in WT and KD HEK293T cells. After 24h, the cells were treated with 250nM Torin1 or DMSO for 2h. **(C)** The methylated and unmethylated reporter mRNAs, as in **Figure 1**, were transcribed *in vitro* and transfected into WT HEK293T cells pre-treated for 2h with 250 nM Torin1 or DMSO. After transfection, the cells were additionally treated with Torin1 or DMSO for 2 h. NlucP activity in Torin1-treated cells was normalized to that in DMSO-treated cells. Values are the means of at least three independent replicates. The error bars show standard deviations. A two-sided Student’s t-test was used to estimate the statistical significance between WT and KD cells **(B)** or between methylated and unmethylated mRNAs **(C)**. ***p<0.001, NS – non-significant. RLU: relative luciferase units.

For comparison, identical plasmids were transfected into the wild-type (WT) HEK293T cells and cells with knockdown of m^6^A methyltransferases *METTL3* and *METTL14* (KD). In KD cells, the synthesized reporter mRNAs were hypomethylated due to the reduced methyltransferase activity (**Figure 2A**).

The translation of *RPL32* reporter mRNA, a canonical mTOR target, was significantly reduced upon mTOR inhibition, and the same effect of lower magnitude was observed for *HSP70*. (**Figure 2B**). In turn, translation levels in KD cells did not differ from WT. However, *SLU7* mRNA translation efficiency was insensitive to mTOR inhibition in wild-type HEK293T cells, but was reduced upon mTOR inhibition to mTOR inhibition in KD cells. This is consistent with the quantitative data on m^6^A methylation^47,48^, which show that *SLU7* mRNA has a higher methylation level than that of *RPL32*. Thus, we hypothesize that m^6^A methylation of *SLU7* mRNA makes it resistant to translation inhibition in WT, and this effect diminishes when the activity of the methyltransferase complex is reduced (KD).

To prove the direct contribution of m^6^A to the translation of reporter mRNAs, we used to *in vitro-synthesize* unmethylated and m^6^A-methylated mRNAs for mRNA transfection. By doing so, we confirmed that methylated mRNAs are less sensitive to mTOR inhibition in HEK293T cells (**Figure 2C**), consistent with the results obtained with CFTS (**Figure 1A,B**).

All in all, we conclude that m^6^A methylation along the mRNA reduces its sensitivity to cap-dependent translation inhibition, and the sensitivity of particular mRNAs depends on their level of m^6^A methylation.

### Transcriptome-wide changes in the translation efficiency upon Torin treatment depend on methylation

Above, we have shown that the resistance of particular mRNA to mTOR inhibition hinges on the presence of an m^6^A modification. To explore this phenomenon at a larger scale, we carried out Ribo-Seq and RNA-Seq analyses in WT HEK293T and HEK293T with *METTL3/14* knockdown in normal conditions and upon the mTOR inhibition.

Comparing relative ribosome occupancy (mTOR inhibition normalized to normal conditions), we observed that the translation inhibition from Torin1 was more pronounced in the KD than in the WT cells. This is clearly illustrated in **Figure 3A** as ∼70% of transcripts fell below the diagonal, which agrees with the assessment of the total translation levels and the mTOR pathway activity (**Figure S3**). Consistently, the total level of protein synthesis, measured by azidohomoalanine incorporation, was approximately 40% lower in the KD cells compared to the WT under normal conditions.

**Figure 3.**
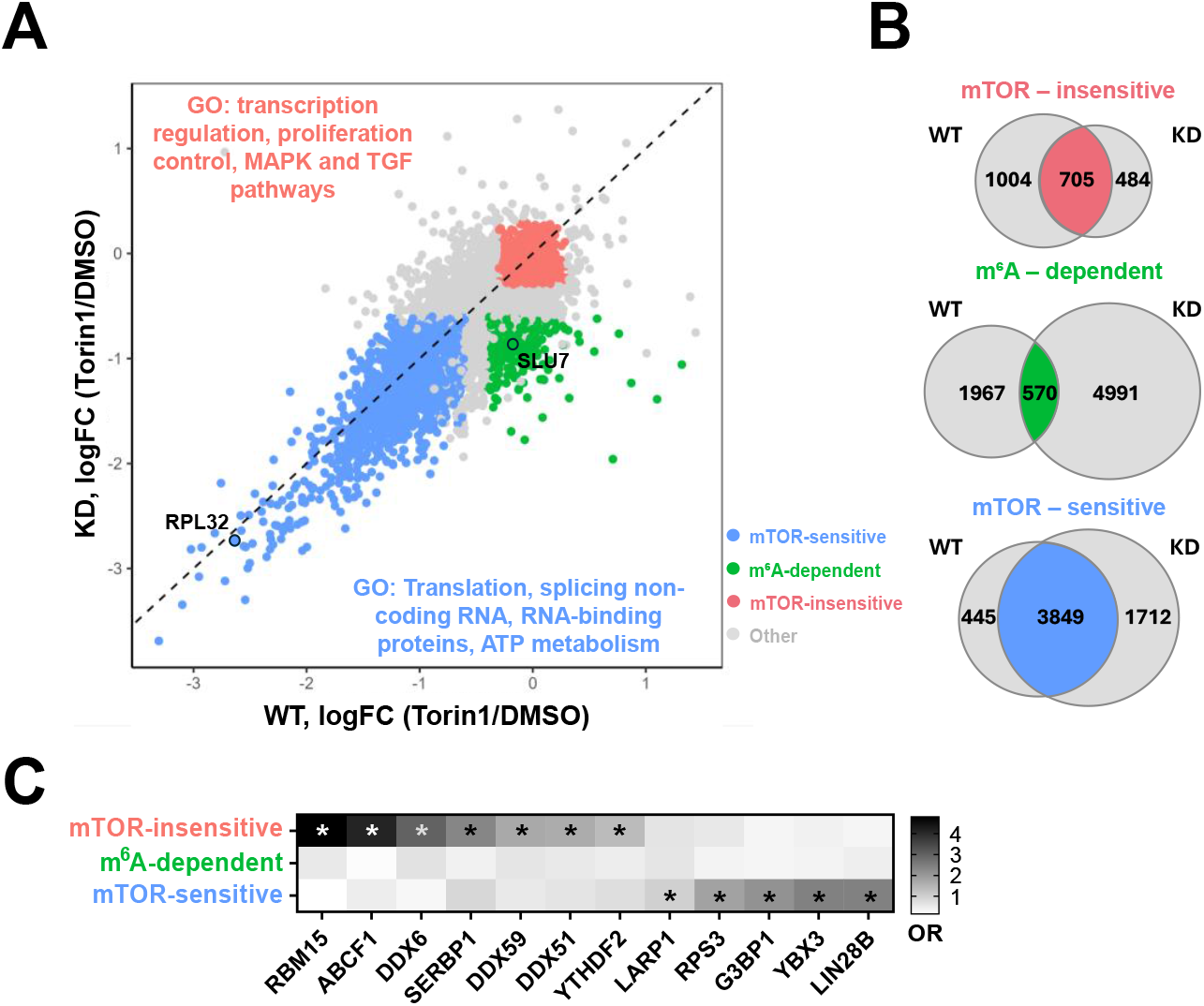
Comparison of translation efficiency in *METTL3/14* knockdown (KD) and wild-type (WT) cells under mTOR inhibition. **(A)** Scatter plot comparing translation changes (Torin1/ DMSO) in KD relative to WT. A diagonal line (x=y) is shown for reference. The values are log_2_-fold changes (Torin1 to DMSO) of the ribosome footprint counts (Ribo-Seq) normalized to the overall transcript abundance (RNA-Seq). Gene groups are colored according to the gene sets depending on the sensitivity to the mTOR inhibition. **(B**) Euler diagrams illustrating the overlap between KD and WT in each gene group. **(C)** RBP targets enrichment. * FDR < 0.05, OR: odds ratio.

Following the Torin1 treatment, the global translation was further reduced by 3.5-fold in the KD cells and by 2.5-fold in the WT cells. Western blot analysis revealed the decreased levels of eIF4E, AKT, and the mTOR kinases in the KD cells (**Figure S3A**), suggesting that the reduced expression of methyltransferases may affect mTOR signaling directly. This observation aligns with an earlier study^12^, which reported that the *METTL3* knockdown amplifies the overall translational sensitivity to mTOR inhibitors.

From Ribo-Seq data, we identified the groups of transcripts that significantly differed in their response to mTOR inhibition (**Figure 3A**). The mTOR-insensitive group (in red) comprised mRNAs whose translation did not change upon the mTOR inhibition in both the WT and the KD cells. Gene Ontology analysis revealed that this group is enriched in transcripts involved in transcriptional regulation, cell proliferation, and MAPK and TGF-β signaling pathways (**Figure S2C**). Using eCLIP, PAR-CLIP, and CLIP-Seq data^42–44,46,53^, we additionally performed enrichment analysis for mRNA targets of RNA-binding proteins (RBPs). By doing so, in this group we detected the enrichment of targets for several RBPs, which might be involved in translation control, particularly in promoting cap-independent initiation (**Figure 3C**). Among those, there were three helicases, namely, DDX6, DDX51, and DDX59. Of those, DDX6 interacts with YB-1 in the 3′ UTR of target mRNAs, promoting eIF4E recruitment and thereby stimulating translation through a non-canonical mechanism^54^. Another notable example is the ribosome-associated protein SERBP1 that has been observed in both actively translating polysomes^55^ and in inactive 80S complexes where it may block the mRNA channel^56^, although its precise function in translation remains to be elucidated.

Next, we focused on the mTOR-sensitive group of mRNAs (**Figure 3A,B**, in blue), which comprised the transcripts with reduced translation under mTOR inhibition in both WT and KD cells. This group was enriched with mRNAs encoding RNA-binding proteins, ribosomal proteins, and splicing factors. Many of these transcripts (including *RPL32*, in agreement with the reporter experiments) contain 5′ TOP motifs (**Table S3**), which are known to mediate sensitivity to mTOR signaling (Meyuhas et al., 2015). RBP targets enrichment analysis revealed several relevant RNA-binding proteins (**Figure 3C**). LARP1 selectively suppresses the translation of TOP-containing mRNAs under mTOR inhibition^57^. LIN28B has been shown to enhance translation of the ribosomal protein mRNAs, well-known mTOR mRNA targets^58^. RPS3 facilitates the translation of TISU-containing transcripts, highly dependent on eIF4E and eIF4G^59–61^. G3BP1 may obstruct cap-dependent initiation by interacting with eIF4E^62^. Finally, YBX3 is known to be involved in translation control and mRNA stability regulation^63^.

Finally, the m^6^A-dependent group (**Figure 3A,B**, in green) included mRNAs whose translation was selectively reduced under mTOR inhibition in KD cells but remained largely unaffected in WT cells. For example, *SLU7* mRNA, in agreement with reporter experiments, showed only a modest decrease in translation (∼13%) upon mTOR inhibition in wild-type (WT) cells, but its translation was suppressed by approximately 50% in KD cells (**Figure 2B**). Thus, for this gene group, the observed reduction in translation is directly related to the reduced level of m^6^A methylation in the KD cells. We did not detect any significant RBP target enrichment for this group of mRNAs (**Figure 2C**).

Overall, we also analyzed the length and GC composition of the transcripts between groups, as these parameters could influence mTOR-dependent regulation of translation^64,65^, but did not detect any notable differences (**Figure S2A**,**B**).

### m^6^A methylome confirms that the translation sensitivity to mTOR inhibition depends on the transcript-wide m^6^A level

With the gene groups defined above (mTOR-sensitive, m^6^A-dependent, mTOR-insensitive), we examined the group-specific fraction of methylated transcripts and transcript-wide methylation levels. To this end, we employed quantitative single-nucleotide resolution m^6^A profiles obtained in HEK293 cells with GLORI^48^ and in HeLa cells with eTAM-Seq^47^. The fraction of transcripts containing at least one methylated position (with any degree of methylation) was comparable between the gene groups (**Figure 4A**).

**Figure 4.**
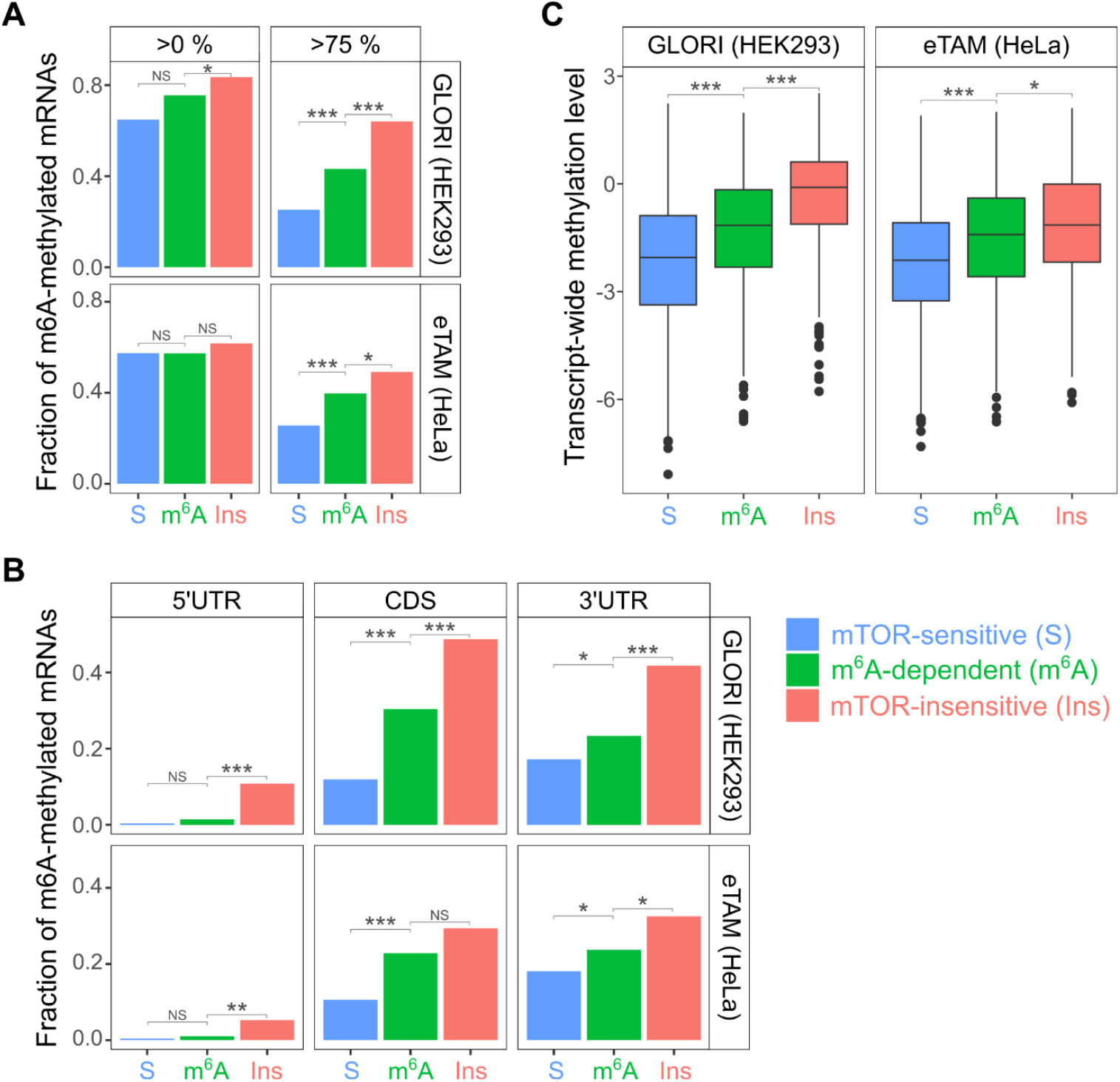
Comparative analysis of m^6^A methylation detected with GLORI (HEK293) and eTAM-Seq (HeLa). **(A)** The fraction of transcripts in each group containing at least one methylated position (>0%) or at least one position methylated over 75%. **(B)** The group-specific fraction of transcripts having at least one position methylated over 75% in a given transcript region (5’ UTR, 3’ UTR, or CDS). **(C)** Normalized total transcript m^6^A methylation, log_2_-scale. * - p < 0.05, ** - p < 10^-3^, *** - p < 10^-5^. A, B: two-tailed Fisher’s exact test, pairwise; C: two-tailed Mann-Whitney U test.

However, if only highly methylated positions (>75% m^6^A) are taken into account, the fraction of methylated transcripts is different across the groups: mTOR-sensitive < m^6^A-dependent < mTOR-insensitive (**Figure 4A**), i.e., the least methylated mRNAs are the most sensitive to mTOR and vice versa. This effect does not depend on the transcript region (**Figure 4B**), although 5’ UTRs rarely include highly methylated positions and thus make only a limited contribution to the total transcript-wide methylation level. This suggests that m^6^A in 5′ UTR alone cannot explain the reduced sensitivity of particular mRNAs to the cap-dependent translation inhibition.

Further, the translation regulation under mTOR inhibition is unlikely to depend on individual m^6^A sites only, but most probably also relies on the overall methylation level along the transcript. To test this hypothesis, we estimated the total m^6^A level along the transcript: the sum of methylation levels of all adenines normalized to the number of adenines (**Figure 4C**). The resulting values showed the same trend as for individual highly-methylated positions: mTOR-sensitive < m^6^A-dependent < mTOR-insensitive.

The gene groups were defined by comparing WT and KD cells, where the reduction in methyltransferase activity decreased the total methylation level. Consequently, in the m^6^A-dependent group, the reduction of the methylation level was sufficient to enable mTOR sensitivity. The genes of the mTOR-insensitive group most likely preserved the total methylation at a level sufficient to resist mTOR inhibition. Thus, there should be a certain threshold level of the total transcript-wide m^6^A methylation, necessary and sufficient to maintain translation under mTOR inhibition.

### The sensitivity to mTOR inhibition saturates at a certain m^6^A methylation level

To test the methylation threshold hypothesis, we transcribed the RPL32 reporter with different ATP to m^6^-ATP ratios (**Figure S4**) and analyzed the translation efficiency of the resulting mRNAs, which differed quantitatively in overall methylation *in vitro* and *ex vivo* upon mTOR inhibition (**Figure 5**).

**Figure 5.**
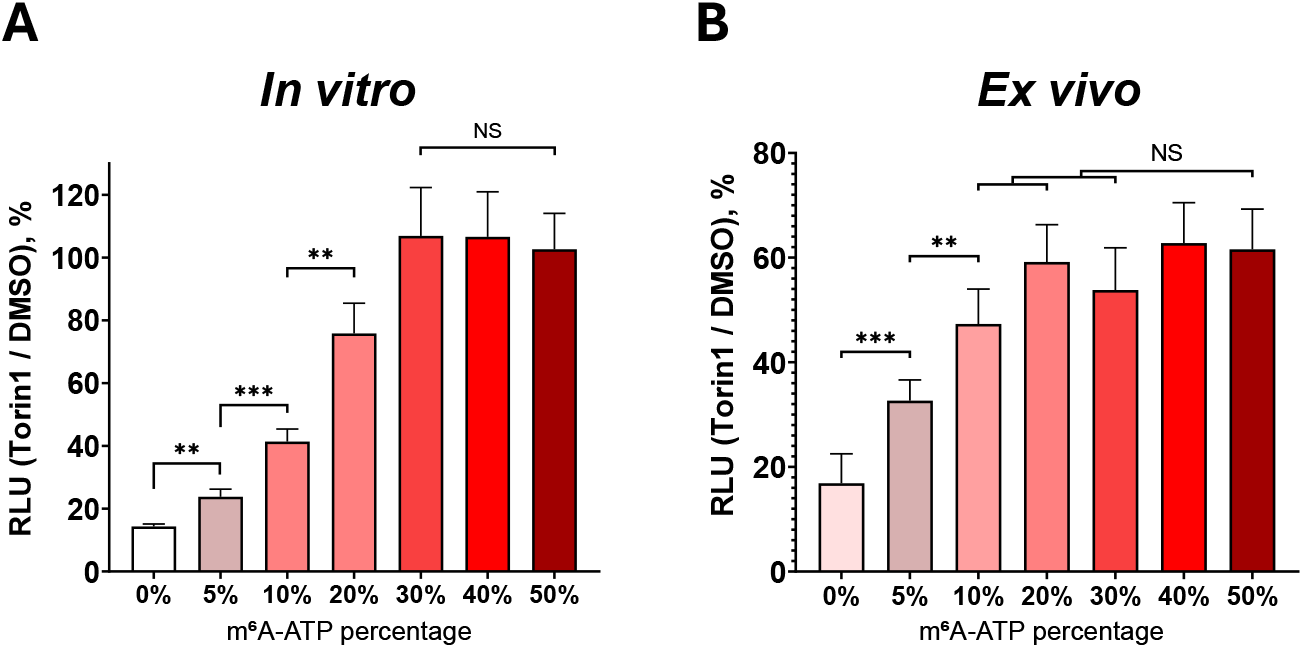
Increasing m^6^A level saturates *RPL32* translation inhibition upon Torin treatment. The barplots show the relative luciferase activity for the *RPL32* reporter mRNAs transcribed *in vitro* with the indicated m^6^ATP percentage (X-axis). **(A)** mRNAs were translated in CFTS prepared from HEK293T cells treated with 250nM Torin1 or DMSO. NlucP activity in CFTS obtained from Torin1-treated cells was normalized to that in the CFTS from DMSO-treated cells. **(B)** mRNA transfected into WT HEK293T cells, pre-treated for 2h by 250nM Torin1 or DMSO. The mRNAs were transfected together with furimazine, and luciferase activity was measured in living cells at 40 min for the CFTS assay and at 80 min for the ex vivo assay. NlucP activity in Torin1-treated cells was normalized to that in DMSO-treated cells. Values are the means of at least three independent replicates. The error bars show standard deviations. The results were analyzed using the two-tailed Student’s t-test, with the significance levels indicated as follows: *p < 0.05; **p < 0.01; ***p < 0.001.

This experiment revealed that m^6^A methylation reduces sensitivity to mTOR inhibition quantitatively. Importantly, (1) the effect is exhibited even at the methylation level as low as 5%, and (2) the effect saturates at 20–30% methylation (**Figure 5A, B**).

## DISCUSSION

In this work, we demonstrated that the m^6^A modification of mRNA plays a key role in the regulation of translation during the inhibition of cap-dependent initiation, specifically, upon inhibition of the mTOR pathway. We showed that m^6^A preserves translation initiation efficiency under stress conditions, probably by promoting the alternative initiation mechanism. Furthermore, we found that the level of m^6^A quantitatively determines the mRNA translation resistance to the mTOR inhibition, and a certain threshold level of the m^6^A modification is sufficient to maintain efficient translation.

The accepted consensus was that m^6^A methylation of mRNA influences translation depending on the location of the modification within the transcript. A major role is attributed to m^6^A sites located in the 5’ UTR, which enhance the recruitment of the cap-independent 43S preinitiation^11,12^. Additionally, m^6^A modifications in the 3′ UTR or near the stop codon can also promote cap-independent initiation through eIF3 recruitment and mRNA circularization^16–18^. On the other hand, m^6^A modifications in the coding sequence (CDS) are believed to influence the translation elongation rate rather than initiation^19,20^.

Contrary to the accepted view, here we showed that the cap-independent translation initiation depends on the m^6^A distributed along the entire mRNA length. This indicates the importance of m^6^A for maintaining mRNA translation initiation despite its negative impact in the elongation phase^21^. Moreover, recent studies have shown that m^6^A in the 5′ UTR is insufficient for driving translation under stress conditions^15^. Also, m^6^A sites in the 5’ UTR are less frequent compared to those in the CDS and 3’ UTR, and they tend to be methylated at lower levels^47,48^. Thus, we consider the impact of m^6^A sites along the transcript on translation initiation to be seriously underestimated.

In *METTL3/14* knockdown cells, the translation sensitivity to mTOR inhibition was non-uniform for different mRNAs. Specifically, the m^6^A-dependent group exhibited reduced ribosome occupancy, while the mTOR-insensitive group maintained translation levels comparable to those observed in WT cells. The regulation of translation within the mTOR-insensitive group may involve mechanisms independent of the m^6^A-mediated control. An explanation can be found in the enrichment of targets of particular RBPs, such as DDX6, which are known to promote translation through the eIF4E recruitment independently of m^6^A modifications (**Figure 3C, Table S3**). Some mTOR-insensitive mRNAs might be in fact m^6^A-methylated in KD cells, as a notable fraction of m^6^A sites remains methylated upon *METTL3* or *METTL14* knockdown^48^. A linked explanation is the certain threshold level of the m^6^A methylation (**Figures 4-5**), sufficient to resist the mTOR inhibition. Indeed, the mTOR-insensitive group is enriched in targets of m^6^A readers, as well as RBM15, a component of the methyltransferase complex. These RBPs may collectively contribute to cap-independent initiation under conditions of mTOR inhibition.

Initiation factor binding assays revealed that eIF4A, eIF4G, and eIF4E have stronger affinity for the m^6^A-modified mRNAs compared to unmethylated controls, even in the presence of Torin1 (**Figure 1C**). This suggests that the mechanism underlying the resistance of methylated mRNA to the inhibition of the cap-dependent initiation is realized through enhanced affinity of the eIF4F complex components. The recruitment of eIF4F components may occur through binding to eIF3 or ABCF1^11,12^ or through the interactions mediated by YTHDF3 with DAP5^66^.

All in all, our findings highlight the importance of transcript-wide m^6^A methylation in promoting translation under mTOR inhibition. Maintaining adequate methylation, in conjunction with RBPs and translation initiation factors including eIF4F complex, seems essential for preserving translation under stress conditions. We emphasize the adaptive significance of m^6^A, shifting the focus from its precise positioning to its general presence as a critical regulator of translation.

## ACKNOWLEDGEMENTS

This study was carried out using resources of the Skoltech Genomics Core Facility (Skolkovo Institute of Science and Technology).

## AUTHOR CONTRIBUTIONS

Egor A. Smolin: Conceptualization, Investigation, Formal analysis, Methodology, Validation, Writing—original draft, Visualization; Andrey I. Buyan: Formal analysis, Investigation, Methodology, Software, Writing—original draft, Visualization; Anton A. Buzdin: Methodology, Data curation, Writing—review & editing; Dmitry N. Lyabin: Conceptualization, Methodology, Writing—review & editing, Project administration, Funding acquisition; Ivan V. Kulakovskiy: Methodology, Software, Data curation, Writing—review & editing, Project administration; Irina A. Eliseeva: Conceptualization, Formal analysis, Visualization, Writing— review & editing, Project administration

## SUPPLEMENTARY MATERIALS

Supplementary Data are available online.

## DATA AVAILABILITY

The data are deposited to the NCBI GEO under the accession number GSE309586, GSE309588.

