## Supplementary material for "Transcript-wide m^6^A methylation defines the efficiency of the cap-independent translation initiation": Figure S1A

### Supplementary Figures

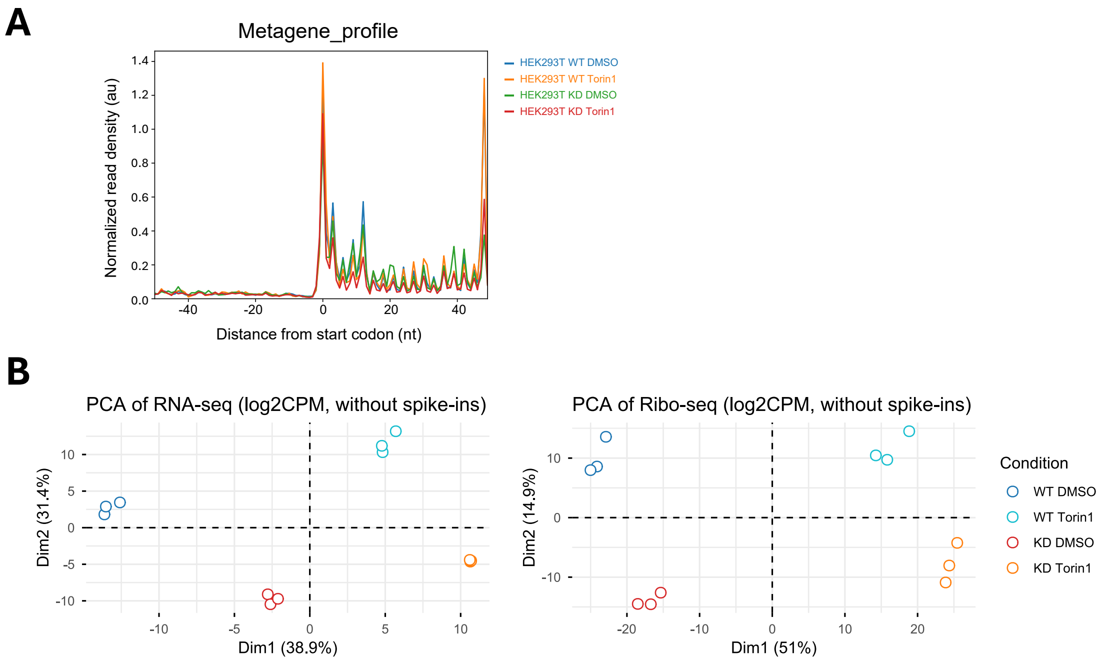


**Supplementary Figure S1. PCA and metagene profiles around start codons.**

**(A)** Metagene profiles representing the P-site position of ribosome footprints are summed up across expressed annotated coding transcripts and anchored at the translation initiation sites. Each profile is normalized to the total counts within a 250 nt [-50;+200] window surrounding the start codons. The plots demonstrate a clear 3-base periodicity within the CDS regions.

**(B)** Principal component analysis (PCA) of normalized RNA-Seq, Ribo-Seq, and RIP-Seq data. The percentage of variation explained by a particular principal component (PC) is indicated in the axis label. Point colors reflect the experiment type.

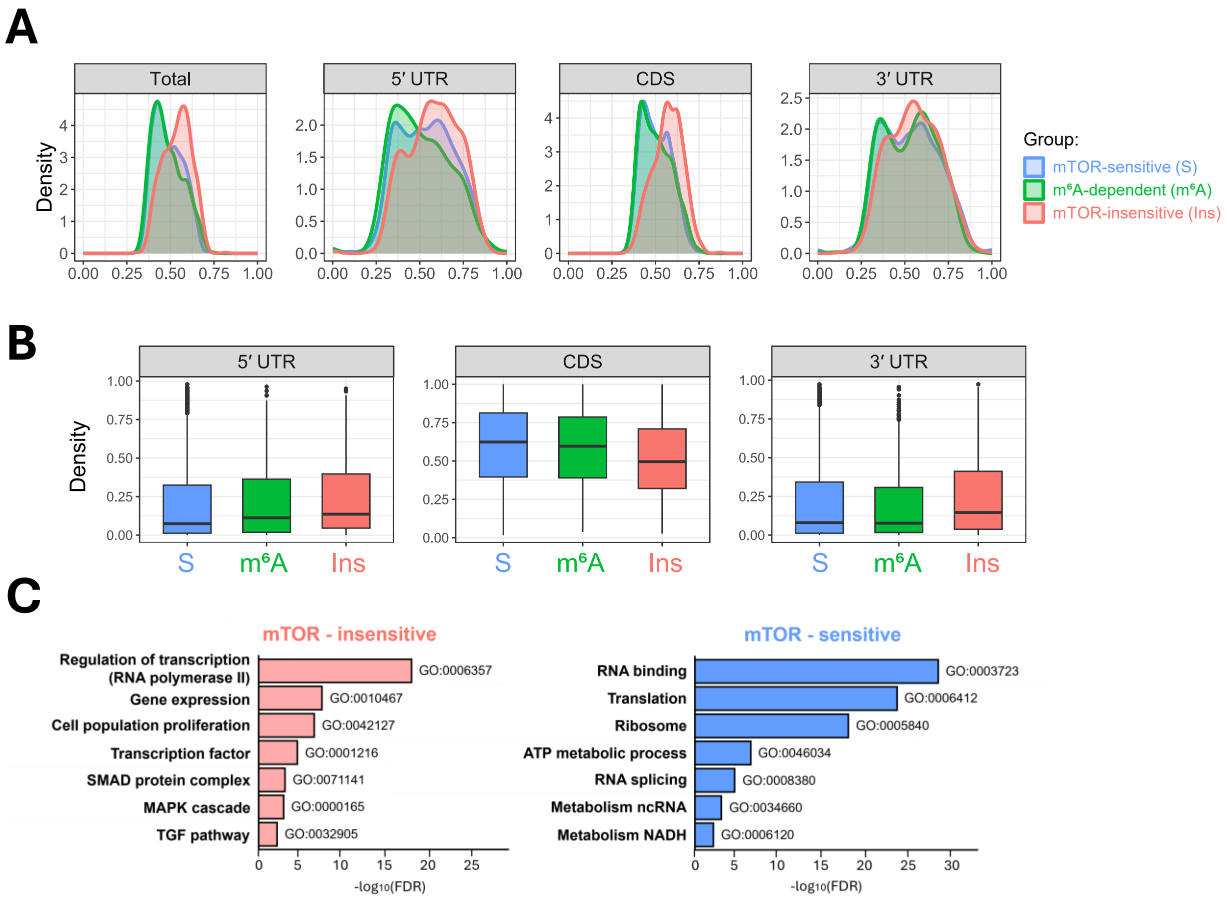


**Supplementary Figure S2. Characteristics of mTOR-sensitive, mTOR-insensitive, and m⁶A-dependent mRNA groups.**

**(A, B)** Distributions of **(A)** transcript length and **(B)** GC-content within different mRNA regions (5′ UTR, CDS, and 3′ UTR) for the indicated mRNA groups: mTOR-sensitive, mTOR-insensitive, and m⁶A-dependent. Length and GC content distributions are shown as box-and-whisker plots.

**(C)** Top 10 significantly enriched Gene Ontology (GO) biological process (BP) terms (FDR < 0.05) identified for each group using GSEA.

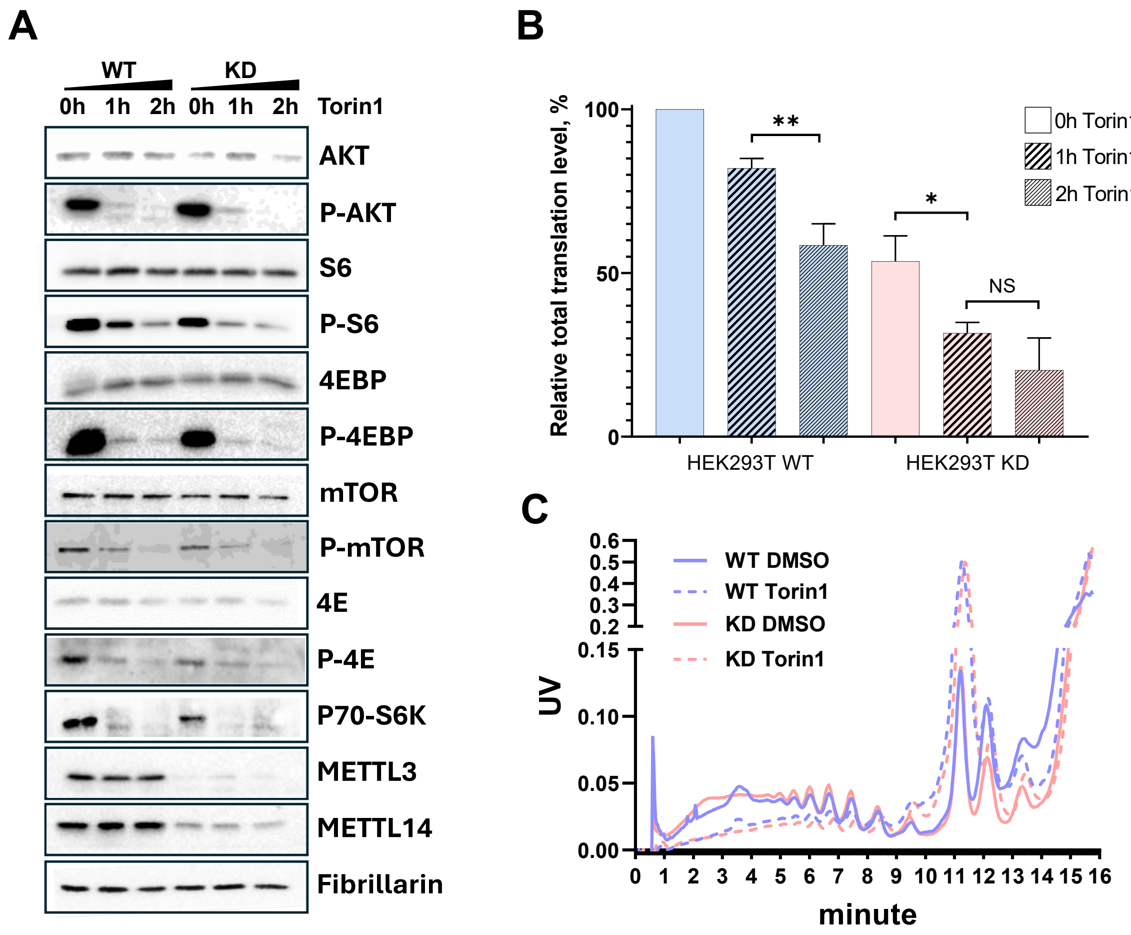


**Supplementary Figure S3. Changes in global translation and major components of the mTOR pathway.**

**(A)** Western blot analysis of HEK293T WT and KD cells under normal and mTOR inhibiting conditions. Cells were treated with 250 nM Torin 1 or DMSO for 1 or 2 hours. Fibrillarin was used as a loading control.

**(B)** Global translation levels in the indicated cells were assessed by azidohomoalanine incorporation followed by a bioorthogonal click reaction with Alexa Fluor 488. Protein fractions were separated by SDS–PAGE, and incorporated fluorescence was visualized using Alexa signal detection. The results were analyzed using a two-tailed Student's *t*-test, with significance levels indicated as follows: *$p$ < 0.05; **$p$ < 0.01.

**(C)** Lysates of HEK293T WT and KD cells were separated by centrifugation through a linear 15–45% sucrose gradient at 45,000 rpm for 1.2 h in an SW-60 rotor.

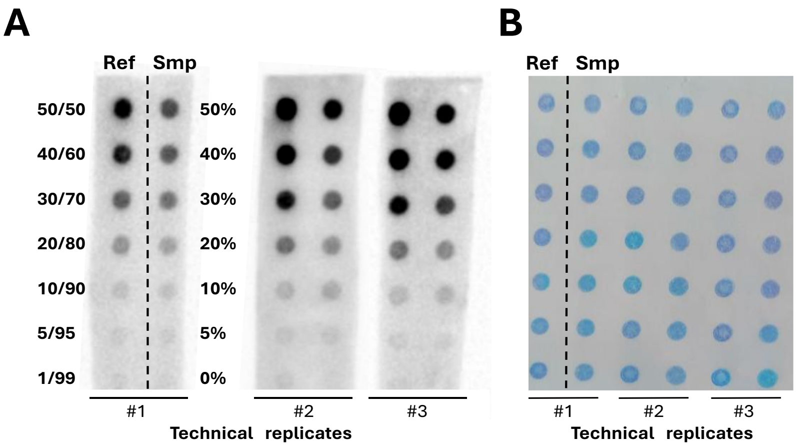


**Supplementary Figure S4. Dot blot analysis of *RPL32* mRNA with defined m⁶A levels.**

The methylation level in reporter mRNAs was controlled by adjusting the ATP/m⁶ATP ratio during in vitro transcription. For dot blot analysis, 500 ng of each mRNA (the percentage of m⁶ATP is indicated as Samples, Smp) was spotted per dot onto a nylon membrane. As a reference (Ref), fully methylated *RPL32* mRNA (100% m⁶ATP) was mixed with unmethylated *RPL32* mRNA (0% ATP) to generate standards corresponding to 1%, 5%, 10%, 20%, 30%, 40%, and 50% m⁶A. (A) Detection using an m⁶A-specific primary antibody followed by HRP-conjugated secondary antibodies. (B) Methylene blue staining of the same membrane as a loading control.

### Supplementary Methods

#### Analysis of mRNA distribution between polysomes and free mRNPs

HEK293T WT and KD cells were grown to 70–80% confluency, cooled on ice, and washed twice with ice-cold PBS containing 0.1 mg/ml cycloheximide. Pre-treatment with cycloheximide was omitted to avoid artificial accumulation of initiation complexes[1]. Cells were lysed directly on the plate in 400 µl of polysome extraction buffer (15 mM HEPES-KOH, pH 7.6, 5 mM $MgCl_2$, 0.3 M NaCl, 1% Triton X-100, 0.1 mg/ml cycloheximide, 1 mM DTT) and incubated on ice for 10 min. Lysates were clarified by centrifugation at 12,000 × g for 10 min at 4°C.

Supernatants (45 µl) were layered onto 4.5 ml of 15–45% sucrose gradients (15 mM HEPES-KOH, pH 7.6, 5 mM $MgCl_2$, 100 mM KCl, 0.1 mM EDTA, 0.01 mg/ml cycloheximide) and centrifuged in an SW-60 rotor (Beckman Coulter) at 45,000 rpm for 70 min at 4°C. Gradients were fractionated from the bottom with continuous monitoring at 254 nm.

#### Measurement of total translation level per cell

Global translation was assessed by azidohomoalanine (AHA) incorporation followed by conjugation with Alexa Fluor 488 alkyne, as described by Presolski et al[2]. Cells grown to 60–70% confluency were incubated for 45 min in methionine-free DMEM (without FBS), then for 2 h in methionine-free DMEM containing 50 µM AHA. After washing with ice-cold PBS, cells were lysed in 1% SDS, 50 mM Tris-HCl (pH 8.0), with 250 U/ml Benzonase.

Lysates (≈200 µg protein) were reacted with Alexa Fluor 488 alkyne in the presence of $CuSO_4$–THPTA, aminoguanidine, and sodium ascorbate for 1 h at room temperature in the dark. Proteins were precipitated with methanol/chloroform, washed, air-dried, and resuspended in 2× SDS–PAGE sample buffer.

Equal protein amounts were separated by SDS–PAGE, and Alexa Fluor 488–labeled proteins were visualized using a ChemiDoc MP imaging system (Bio-Rad, USA) and quantified with Image Lab. Total protein was determined by Coomassie Brilliant Blue G-250 staining and analyzed with OptiQuant. Translation per cell was expressed as Alexa Fluor 488 fluorescence normalized to total protein content.
